# Divergent selection in Mediterranean pine stands on local spatial scales

**DOI:** 10.1101/2023.02.16.528264

**Authors:** Katharina B. Budde, Christian Rellstab, Myriam Heuertz, Felix Gugerli, Miguel Verdú, Juli G. Pausas, Santiago C. González-Martínez

## Abstract

The effects of selection on an organism’s genome are hard to detect on small spatial scales, as gene flow can erase signatures of local adaptation. Most genome scans to detect signatures of environmental selection are performed on large spatial scales, however divergent selection on the local scale (e.g. between contrasting soil conditions) has also been demonstrated, in particular for herbaceous plants. Here we hypothesize that in topographically complex landscapes, microenvironment variability is strong enough to leave a selective footprint in genomes of long-lived organisms. To test this, we investigated paired south- versus north-facing *Pinus pinaster* stands in a Mediterranean mountain area. While north-facing (mesic) stands experience less radiation, south facing (xeric) stands represent especially harsh conditions, particularly during the dry summer season. Outlier detection revealed five putatively adaptive loci out of 4,034, two of which encoded non-synonymous substitutions. Additionally, one locus showed consistent allele frequency differences in all three stand pairs indicating divergent selection despite high gene flow on the local scale. Functional annotation of these candidate genes revealed biological functions related to abiotic stress response in other species. Our study highlights how divergent selection shapes the functional genetic variation within populations of long-lived forest trees on local spatial scales.

## Introduction

Spatially heterogeneous environments exert divergent selection pressures and can contribute to maintaining high levels of adaptive genetic variation within populations. However, understanding under which circumstances selection is acting and especially on which spatial scale divergent it can be detected remains poorly understood. Studying local adaptation in forest tree species is an important endeavor especially under current climate change [1,2]. Numerous studies have already revealed loci potentially involved in environmental adaptation. However, most of these studies have been conducted on regional to continental scales [e.g., 3–6], as gene flow on small spatial scales can blur the migration–selection equilibrium maintaining local adaptation. Divergent selection on the local scale, e.g. to toxic soil conditions, has often been observed in herbaceous plant species [7,8]. There is increasing evidence that plants exhibit adaptive divergence on very small spatial scales, i.e. on scales of tens of meters of distance in herbaceous species and of hundreds of meters of distance in some woody species (reviewed in [9,10]). Recent studies have started to address the factors shaping local adaptation on the microenvironmental scale in long-lived tree species [11–14]. Several tropical tree species show adaptation to microenvironmental conditions [13,14]. *Eperua falcata* (Fabaceae), for example, showed divergent selection between groups of individuals growing in seasonally flooded bottomlands and adjacent groups growing on dry *terra firme* soils [17,18]. Also, Gauzere *et al*. [19] found evidence for divergent selection acting on growth and phenology traits along an altitudinal gradient within natural stands of European beech (*Fagus sylvatica*) despite high gene flow.

Identifying the genes and gene variants that confer local adaptation, i.e. higher fitness to certain environmental conditions, is of great interest in ecology and evolution. The detection and validation of candidate loci potentially under selection, however, remains challenging. Experimental functional validation is not attainable in non-model species, especially in trees with their long generation times. Previous studies showed that many approaches to detect loci under selection can be prone to false positives (e.g., [20,21]) and that the identified genomic signatures of selection might not always be observed in other locations with similar environmental conditions [22–24]. Therefore, combining several analytical approaches is recommended to reduce false positive detection [25]. Additionally, an appropriate sampling design can increase the power to detect loci involved in local adaptation. Especially, a paired design comprising several pairs of sampling sites with contrasting environmental conditions seems promising for the detection also of loci displaying weak signatures of selection [26]. A simulation study by Lotterhos & Whitlock [21] showed that sampling pairs of nearby populations (i.e. at gene flow distance) with contrasting environmental conditions increases the probability of detecting true positive outlier loci compared to gradient or random sampling designs.

In Mediterranean ecosystems, water availability is one of the most important factors driving selection and plant species are typically well adapted to summer dry conditions [27,28]. Still, considerable microenvironmental variation can be observed especially in topographically complex landscapes, such as Mediterranean mountain systems. Equator-facing slopes receive lower solar radiation flux density, leading to lower evapotranspiration rates and lower daily maximum temperatures during summer drought periods, and therefore show a significantly different composition, structure and density of plant communities as compared to slopes facing pole-wards [29–31]. We hypothesize that, in topographically complex Mediterranean forests, microclimate variability is strong enough to leave a selective footprint on long-lived trees. In this study, we used a robust paired sampling design within a natural population of Maritime pine (*Pinus pinaster* Aiton, Pinaceae) to specifically test for genetic signatures of divergent selection between xeric (south-facing slope) and mesic (north-facing slope) conditions.

## Material and methods

### Study species and sample collection

Maritime pine is a monoecious conifer species growing in the western part of the Mediterranean basin and along the Atlantic coast in south-western Europe. It is pollinated and dispersed by wind. Pollen flow is therefore wide-ranging, following highly leptokurtic dispersal kernels with average dispersal distances of 78-174 m and frequent long-distance dispersal events [32]. Gene flow via seeds is more restricted (average of 26.53 m [33]), but post-dispersal processes, such as the Janzen-Connell effect and microenvironmental variation affecting survival at early life stages can substantially increase effective dispersal distances [34].

For this study, we sampled three pairs of *P. pinaster* stands with contrasting microenvironmental conditions in a natural forest near Eslida in Sierra de Espadán, Eastern Spain (Fig. 1). All *P. pinaster* trees belong to a single gene pool [35,36] and the region is characterized by a warm and dry climate. Stand-replacing crown fire events are common and may take place every few years. Under these conditions regeneration is mostly driven by fire events leading to even aged cohorts [37]. We selected one stand pair consisting of one south-facing slope and trees from a nearby shady valley along a (mostly) north-exposed stream (S1/N1) and two pairs of stands (S2/N2 and S3/N3) with south- (dry and warm) and north-facing slopes (more humid and less warm). For simplicity, we will refer to this sampling design as three pairs of south- and north-facing slopes. In each of the six stands, we haphazardly sampled 25 trees with DBH (diameter at breast height) > 16, making a total of 150 trees (Fig. 1, Table 1). All trees were georeferenced using a Garmin Oregon 550t (Garmin, Wichita, USA), height was assessed using a Digital hypsometer Forestor Vertex (Haglöf, Långsele, Sweden) and the DBH was measured. The maximum straight-line distance between sampled trees was ca. 10 km between stand pairs and 820 m between trees within pairs.

**Table 1.** Paired stand sampling of south- (S) and north-facing (N) slopes for *Pinus pinaster* in the Sierra de Espadán (eastern Spain), and genetic diversity estimates based on 5,024 single-nucleotide polymorphisms (SNPs). ID, identifier for each stand (see Fig. 1); Latitude, latitude in decimal degrees; Longitude, longitude in decimal degrees; Aspect, average aspect in degrees; Altitude, altitude in meters above sea level; Height, tree height in meters with standard deviation, *N*_*S*_, number of samples; *H*_E_, expected heterozygosity; SE, standard error; *F*_IS_, fixation index.

| ID | Lat. | Long. | Aspect<br>[°] | Altitude<br>[m a.s.l.] | $N_S$ | Height<br>[m] (SD) | $H_E$ (SE) | $F_{IS}$ |
| --- | --- | --- | --- | --- | --- | --- | --- | --- |
| Pair 1 |  |  |  |  |  |  |  |  |
| S1 | 39.865 | -0.298 | 185 | 632-666 | 25 | 7.300<br>(1.524) | 0.336<br>(0.003) | -0.018 |
| N1 | 39.866 | -0.298 | 297 | 645-737 | 25 | 8.732<br>(1.334) | 0.330<br>(0.003) | 0.008 |
| Pair 2 |  |  |  |  |  |  |  |  |
| S2 | 39.895 | -0.353 | 159 | 719-763 | 25 | 8.716<br>(1.763) | 0.328<br>(0.003) | 0.013 |
| N2 | 39.895 | -0.351 | 35 | 655-730 | 25 | 12.496<br>(1.900) | 0.337<br>(0.003) | -0.004 |
| Pair 3 |  |  |  |  |  |  |  |  |
| S3 | 39.917 | -0.397 | 181 | 655-728 | 25 | 8.716<br>(1.763) | 0.332<br>(0.003) | -0.005 |
| N3 | 39.913 | -0.389 | 340 | 696-731 | 25 | 11.104<br>(1.981) | 0.326<br>(0.003) | 0.005 |

**Fig. 1.**
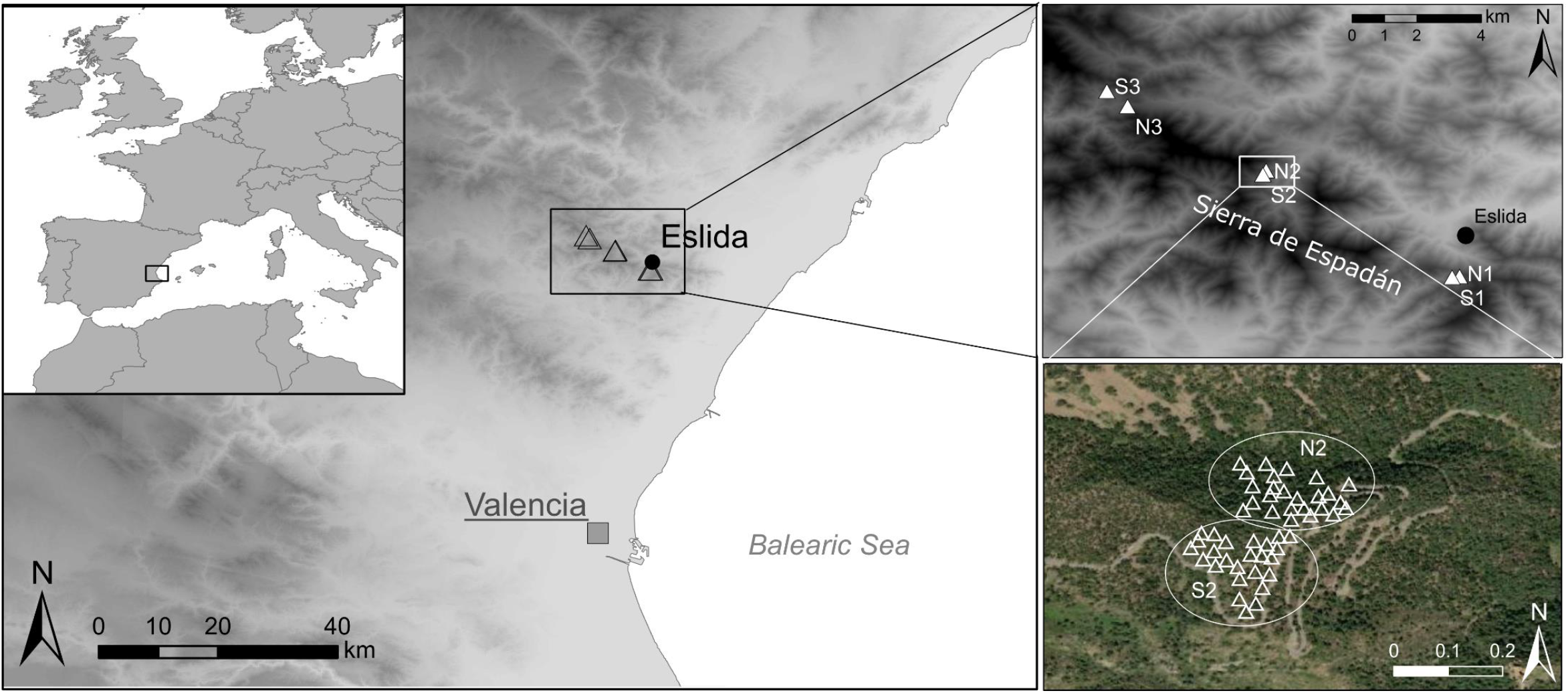
Sample collection of *Pinus pinaster* in three pairs of south- (S) and north-facing (N) slopes in Sierra de Espadán (eastern Spain) and a detailed view of stands N2/S2 (bottom right).

### DNA extraction and genotyping

Needles were collected from the 150 trees and desiccated using silica gel. Genomic DNA was isolated using the Invisorb® DNA Plant HTS 96 Kit/C kit (Invitek GmbH, Berlin, Germany) following the manufacturer’s instructions.

An Illumina Infinium SNP (Single Nucleotide Polymorphism) array (Illumina, Inc., San Diego, USA) developed by Plomion *et al*. [38] was used for genotyping. This array is enriched in SNPs from genes that showed signatures of natural selection in previous studies [27, 28, 29] or differential expression under biotic and abiotic stress [38] in maritime pine, but most of the SNPs represent potentially neutral polymorphisms. After removing SNPs with uncertain scoring based on visual inspection using *GenomeStudio* Genotyping Module v1.0 (Illumina, Inc.) and monomorphic SNPs, we kept 5,024 high-quality SNPs, of which 4,034 had a minor allele frequency (MAF) > 0.1. The amount of missing genotype data per stand was very low (maximum of 1%). This data set has recently been used to characterize the effective population size in Sierra de Espadán, as part of a meta-study [41].

### Data analyses

First, we characterized the study stands based on the sampled trees’ height and DBH and tested if these phenotypic traits differed significantly between south- and north-facing slopes using a two sample Student’s *t* test on each stand pair run in R v. 4.1.2 [42]. Then, based on the SNP data, we estimated genetic diversity parameters such as observed and expected heterozygosity and the fixation index using the R package *hierfstat* [43]. After this, we tested whether we could detect significant neutral genetic differentiation between the sampled stands by estimating pairwise *F*_ST_ [44] using the complete SNP data set and comparing with neutral expectations from 1,000 permutations. To visualize the neutral population genetic structure inherent to our data, we also performed a Principal Component Analysis (PCA) using the function *dudi*.*pca* implemented in the R package *ade4* [45] and a supervised (i.e. defining each stand as a group) Discriminant Analysis of Principal Components (DAPC) using the *dapc* function in the R package *adegenet* [46] based on all SNP markers. Additionally, we assessed the fine-scale spatial genetic structure (SGS) within each of the three pairs. First, we estimated the pairwise Loiselle kinship coefficient [47] in SPAGeDi v. 1.5d [48] between individuals. The average kinship coefficient per distance class was regressed against the logarithm of spatial distances and significance was assessed based on 10,000 permutations of individual locations. The strength of SGS was estimated as Sp =–b/(1 – F_1_), where b is the regression slope and F_1_ is the average kinship coefficient in the first distance class [49].

To detect loci potentially under selection in slopes with contrasting aspects (south/north) in a hierarchically structured population [50], we used two hierarchical *F*_ST_ outlier detection approaches that take into account the paired sampling design, one implemented in Arlequin v 3.5.2 [51] and the other in BayeScanH, which is especially suitable for small sample sizes [52]. For this, we first defined the pairs and then the aspect of the slopes (south/north) within pairs. In Arlequin *F*_ST_ values can be slightly negative especially on small spatial scale but including loci with negative *F*_ST_ values impedes outlier analyses. Therefore, only SNPs with positive values of estimated *F*_ST_ (1,810 SNPs) were considered in Arlequin analysis (200,000 simulations). We report *F*_SC_ outlier loci for divergence between sites within pairs. To identify outlier loci with BayeScanH, we used the full dataset of 4,034 SNPs with MAF > 0.1, and default parameters with an odds prior of 10. We tested two models, one with the same selection pressure acting between contrasting slopes in the three stand pairs and another one with three independent selection pressures on contrasting slopes within the three pairs. Finally, using the R-script ‘paired_GEA.R’ from https://gitlabext.wsl.ch/rellstab/genotype-environment-associations, we tested if any of the candidate SNPs identified with Arlequin or BayeScanH showed consistent patterns in population allele frequencies between the paired stands in all replicates. For this, we checked whether the differences in population allele frequency had the same sign in all pairs, i.e. whether the population allele frequency in all north-facing slopes was consistently lower or higher than the allele frequency in all south-facing slopes (i.e. the strict sign test). Then, we ran a linear mixed model using the function *lme* implemented in the package *nlme* [53], with population allele frequency as response variable, slope aspect (south/north) as fixed effect and pair as random factor.

The sequences flanking SNPs identified as loci potentially under selection, and associated annotation, were retrieved from Plomion *et al*. [38]. These sequences were newly blasted against the NCBI nucleotide database to check for new functional annotations.

## Results

Tree height was consistently lower in south-than in north-facing slopes and the difference was significant in two out of the three stand pairs (Supp. Mat. Fig. S1.1, Table S1.1), while no significant difference was detected for DBH in any stand pair.

Expected and observed heterozygosity (not shown) were very similar in all six study stands with values around 0.33, resulting in fixation indices close to zero (Table 1). Pairwise genetic differentiation between stands based on all 5,024 SNP markers was weak, ranging from 0.004 to 0.033, but highly significant above zero, with all *P*-values < 0.001 (Supplementary Material, Table S2.1). The DAPC clearly depicted the hierarchical population structure due to the paired sampling design, with stronger genetic differentiation among than within stand pairs (i.e. between south- and north-facing stands; Fig. 2). The hierarchical population structure was also visible but less evident in the PCA plot (Supplementary Material, Fig. S2.1). SGS was significant, showing isolation by distance, in all stand pairs and strongest in pair N3/S3 (Supplementary Material, Fig. S2.2).

**Fig. 2.**
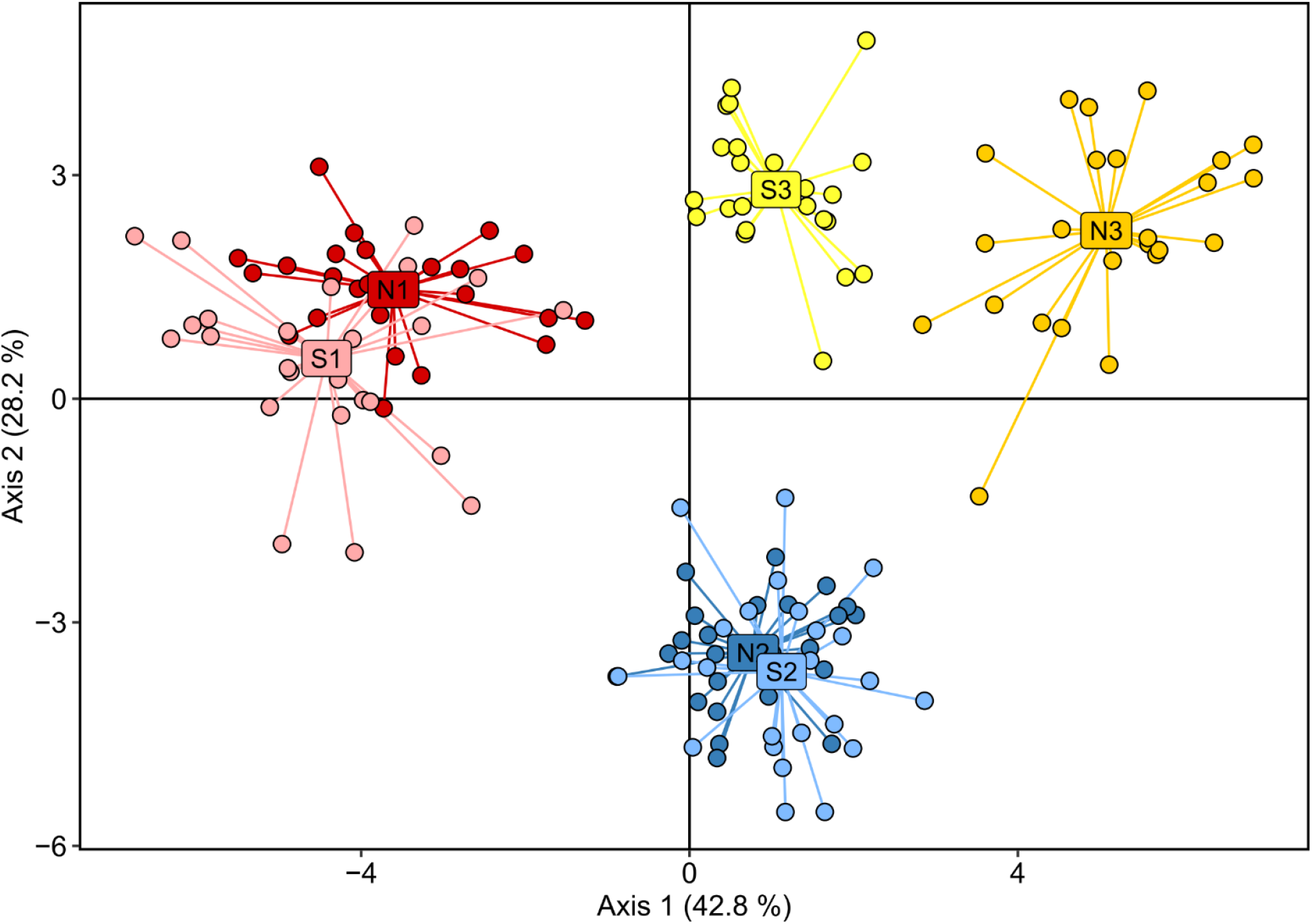
Discriminant Analysis of Principal Components (DAPC) of *Pinus pinaster* samples from Sierra de Espadán (eastern Spain) including three pairs of south- (S) and north-facing (N) slopes based on 5,024 single-nucleotide polymorphisms (SNPs). Each stand is depicted with a different colour and the stand centroid is labelled with the site identifier (see Fig. 1).

In total, 18 SNPs were located above the 99% confidence intervals using the hierarchical island model in Arlequin and, thus, were considered as significant outliers for genetic differentiation between south- and north-facing slopes (Fig. 3, Supplementary Material Table S3.1). Additionally, ten loci were identified as significant *F*_ST_ outliers by BayeScanH when assuming independent selection pressures for each of the three pairs of stands (Supplementary Material, Table S3.1). None of these outlier loci was significant in all three sampling pairs in BayeScanH. Moreover, no significant outlier locus was detected when assuming the same selection pressure in all three pairs of stands.

**Fig. 3.**
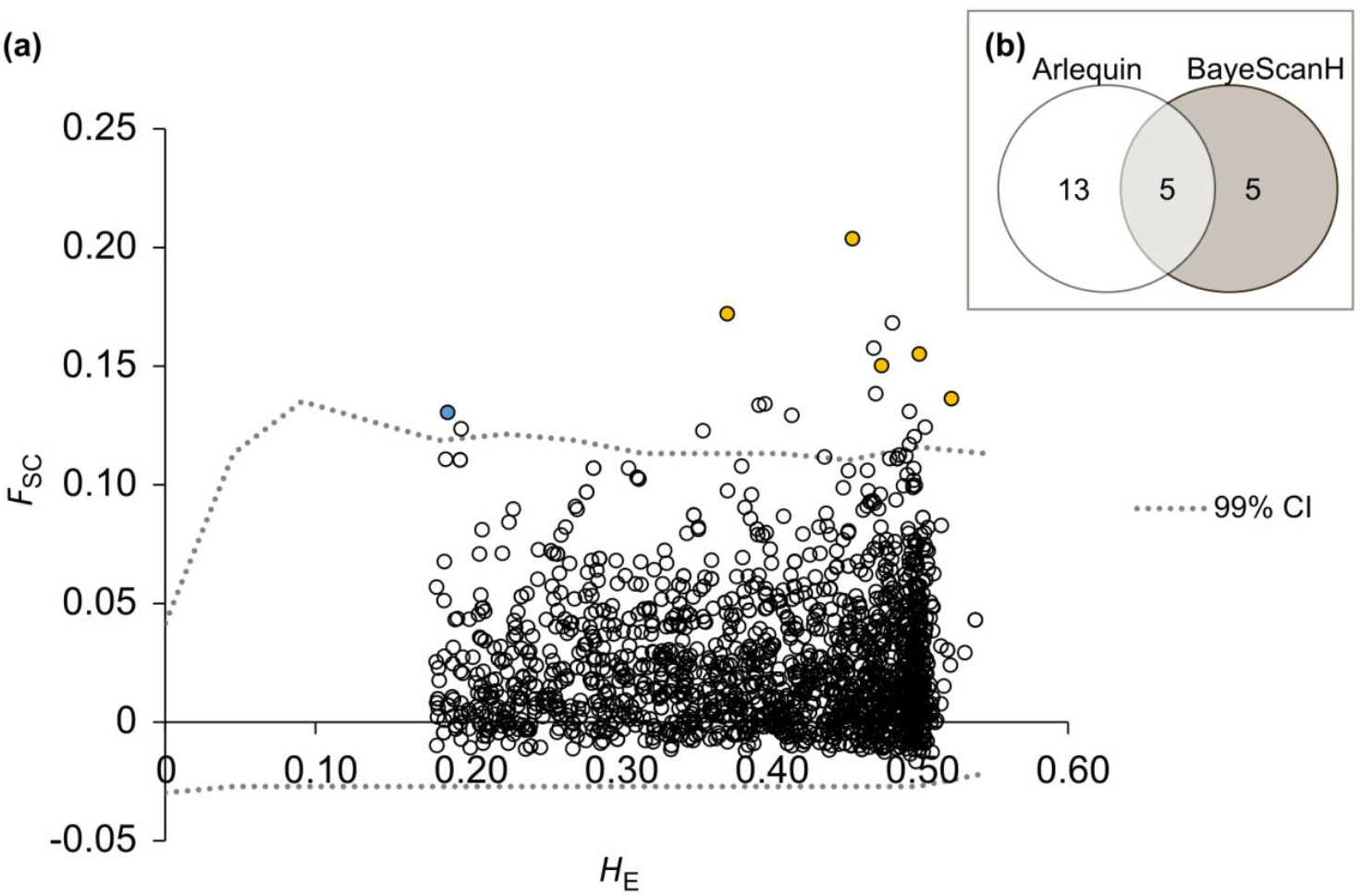
Detection of outlier single-nucleotide polymorphisms (SNPs) using the hierarchical island model (south-vs. north-facing slopes) implemented in Arlequin. (a) *F*_SC_: estimates of locus-specific genetic divergence between stands within pairs; *H*_*E*_: heterozygosity per locus. Dashed lines indicate upper 99% confidence intervals for variation in neutral *F*_SC_ as a function of *H*_E_, indicative of divergent selection. Only AL751008-691 (in blue) showed a consistent shift in allele frequencies in all pairs of stands as indicated by the sign test. Another five loci (in yellow) were also detected as outliers by BayeScanH. (b) Venn diagram showing the overlap of significant outlier loci detected by Arlequin and BayeScanH, respectively.

When comparing the two methods, five loci were identified as outliers by both Arlequin and BayeScanH, and only one additional outlier locus, AL751008_691 detected by Arlequin, showed consistent allele frequency differences between south- and north-facing slopes (Figure 4) and a significant effect of the site aspect as indicated by the linear mixed model (*P*_site type_ = 0.0021). Two out of these six outliers SNPs showed non-synonymous changes and coded for a putative RNA-binding protein and a V-type proton ATPase catalytic subunit, respectively (Table 2).

**Table 2.** Functional annotation of five single-nucleotide polymorphisms (SNPs) detected as significant *F*_ST_ outliers by both Arlequin and BayeScanH and one SNP (AL751008_691) detected only by Arlequin that showed consistent allele frequency patterns in the three stand pairs. The information was retrieved from Plomion *et al*. [38] and confirmed with a new Blast search; non-syn., non-synonymous; leu, leucine; pro, proline; glx, glutamine. NA, not available.

| SNP ID | Polym. | Site type | Protein change | Putative function | Linkage group |
| --- | --- | --- | --- | --- | --- |
| AL751008-691 | [T/C] | NA |  | unknown | LG2 |
| BX250086-1490 | [T/C] | non-syn. | leu → pro | oligouridylate binding protein-like | NA |
| BX251523-1352 | [A/C] | non-syn. | glx → pro | V-type proton ATPase catalytic subunit A | LG10 |
| BX252256_232 | [C/G] | NA |  | Yos1 domain containing protein | NA |
| CT574789-692 | [A/C] | NA |  | unknown | LG12 |
| i09683s215pg | [A/G] | non-coding |  | raffinose_syn domain-containing protein | LG12 |

**Fig. 4.**
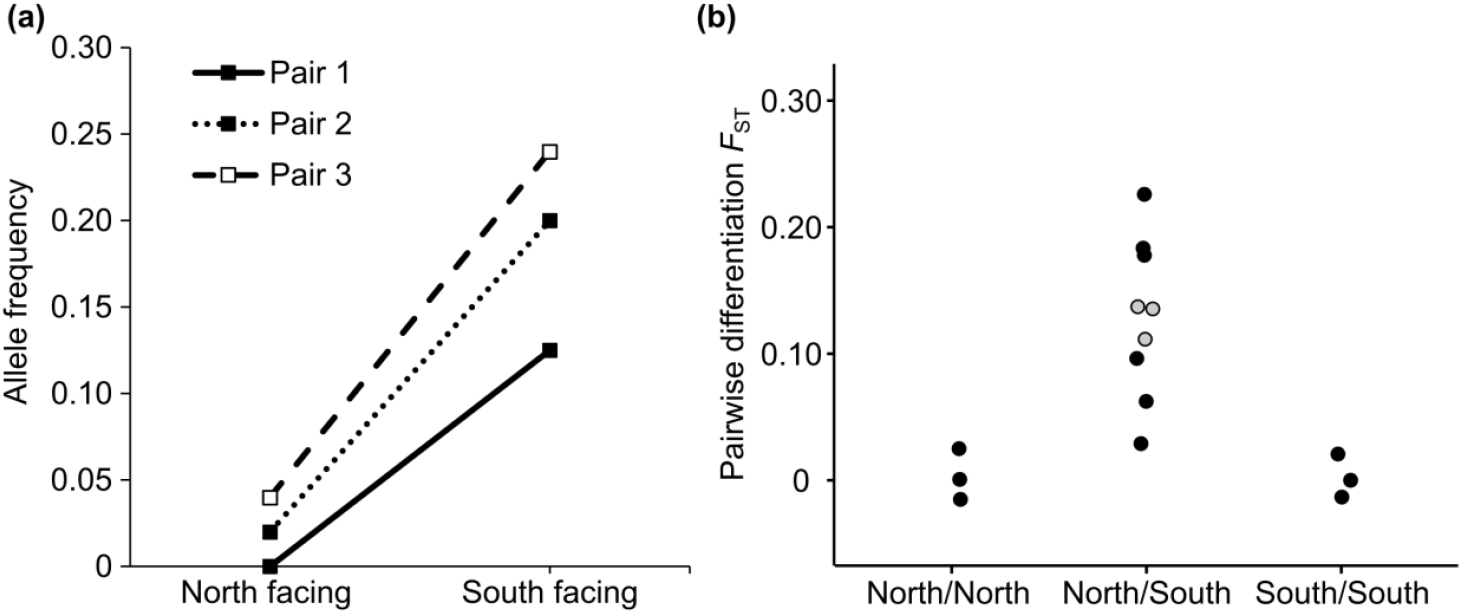
Differences in allele frequencies and genetic differentiation between stand pairs with contrasting aspects. (a) Plot of locus AL751008_691 showing consistent differences in allele frequency between south- and north-facing slopes for all three stand pairs. (b) Pairwise *F*_ST_ between stands, with values between south- and north facing slopes in each sampling location plotted in grey.

## Discussion

The paired sampling design in Sierra de Espadán, contrasting south- and north-facing slopes within a large and continuous *P. pinaster* population, was specifically used to test for microenvironmental adaptation driven by water availability. Paired sampling in stands with contrasting environments, such as dry vs. humid patches, represents a powerful approach to reveal loci under selection [21,25], because it maximizes potential for divergent selection while minimizing the effect of confounding population structure. Several studies successfully employed the paired sampling design to detect loci under selection (e.g. [14,18,54–58]). In conifers, four previous studies revealed loci significantly associated to altitudinal or other microenvironmental gradients in *Abies alba* [57,58], *Pinus halepensis* [56], and *P. pinaster* and *Cedrus atlantica* [14], but only few loci showed consistent patterns of allele frequency shifts along the replicated stand pairs. Here, we specifically tested for consistent patterns of divergent selection on the local scale, with trees growing in direct vicinity, between mesic and xeric stands.

We first showed a hierarchical population genetic structure despite high gene flow in *P. pinaster* within one large population in Sierra de Espadán (eastern Spain). From previous work, it is known that this forest constitutes a single gene pool [35,36]. Moreover, fine-scale spatial genetic structure within continuous populations typically is weak in this wind-pollinated and wind-dispersed species [59]. Therefore, it is remarkable to find significant differentiation among all sampling sites, even between the neighbouring south- and north-facing slopes, clearly depicting the hierarchical structure. This pattern could be caused by phenological differences in flowering time restricting effective gene flow between contrasting slopes due to temporal separation [60]. However, it could also reflect isolation by distance to some degree as indicated by significant SGS (Supplementary Material Fig.2.2). Indeed, differentiation between neighbouring slopes and SGS were strongest for pair N3/S3, which was also the pair with the biggest geographic distance between stands. However, differentiation between directly neighbouring south- and north-facing slopes could additionally be driven by isolation by environment (IBE), which would imply selection against maladapted immigrants resulting in genome-wide patterns of genetic differentiation [61,62]. Furthermore, clonal common gardens of range-wide *P. pinaster* populations show strong local adaptation to different environmental conditions [35,63–65]. In particular, tree height was shown to correlate negatively with maximum summer temperatures (i.e. trees from populations originating in hotter environments tend to be smaller when grown in the same environment), indicating an adaptive response to hot summer temperatures [64]. In forest trees, steep equator-facing slopes usually limit growth, while taller trees are typically found on less steep and pole-facing slopes [66]. In agreement with this, *P. pinaster* trees on south-facing slopes in the Sierra de Espadán tended to be smaller, which could indicate that the populations responded to the harsher environmental conditions either through plasticity or local adaptation.

Second, the complementary approaches to detect outlier loci in hierarchical sampling designs identified five out of 4,034 SNPs (with MAF>0.1) as putatively under divergent selection on local spatial scales. One additional locus (AL751008-691) showed a consistent allele frequency pattern in accordance with microenvironmental adaptation in all three stand pairs. Environmental conditions on south- and north-facing slopes are known to differ strongly, e.g. in light and water availability [67]. Slopes with different aspect are often characterized by differences in composition, structure and density of plant communities [29–31]. Tree species have developed diverse adaptations in response to strong selection pressures in dry environmental conditions [68]. *Pinus pinaster* stands as a suitable study species to test for divergent selection on the local scale. Multisite clonal common gardens comprising range-wide populations already revealed that the species is susceptible to drought. Survival was lowest in the common garden sites with the harshest (dry and hot) conditions [63], and certain alleles at candidate loci associated with climate were connected to a higher probability of survival [35]. Here, we showed that contrasting environmental conditions on different slopes, in direct vicinity and in the presence of gene flow, can also shape the distribution of genetic variation in long-lived forest trees such as *P. pinaster*.

Outlier loci related to differences in drought intensity and temperature have been found in different pine species on range-wide spatial scales. For example, Eckert *et al*. [69], found five outlier loci associated with aridity in *Pinus taeda*. In natural *Pinus albicaulis* populations, Lind *et al*. [70] also identified water availability as a strong driver of genomic adaptation signatures. They detected allele frequency changes at candidate genes along a precipitation gradient on the regional scale in the Lake Tahoe Basin, an ecosystem similar to that studied here (i.e. Mediterranean-type mountains). Candidate gene approaches in maritime pine also found various outlier loci related to drought response and precipitation on large spatial scales [35,39,71] and between shady and sunny stands at the microenvironmental scale [14]. Our study detected a small number of outlier loci potentially related to water availability in maritime pine on the local scale, i.e. within gene flow distance. One of these outlier loci (CT384-490, coding for a non-synonymous change) has been previously associated to winter precipitation on the range-wide scale [35]. Four of the six candidate SNP loci showing strong evidence of local adaptation on small spatial scales were functionally annotated and two of them coded for non-synonymous changes. Locus BX250086 coded for an oligouridylate binding protein-like protein and BX251523 for a V-type proton ATPase catalytic subunit. Locus i09683s215pg, which is coding for a non-synonymous change, is located in a gene encoding for a raffinose_syn domain containing protein. Genes annotated with similar functions have been described to be involved in abiotic stress response, such as drought stress, in other plant species [72–74].

In the last years, reference genomes, even for conifer species with extremely large genomes (> 18 Gbp), have been published [75–77], however, the functional annotation of conifer genomes is still limited and a reference genome for *P. pinaster* is lacking. In this study, we were able to retrieve putative annotations for only four out of six candidate genes, highlighting the need to complete and improve our knowledge of conifer genomes and their functional annotation. In addition, although we were able to identify some candidate loci under divergent selection on the local scale, only one locus showed consistent differences in allele frequencies in all three stand pairs. This is in agreement with a recent study by Scotti et al. [14] where only a small proportion of outlier loci (0.1-1% of all loci depending on the species) showed consistent allele frequency differences between pairs of sites with contrasting conditions indicating that common signatures of selection are scarce. In BayeScanH, significant results were only obtained when assuming three independent selection pressures, which suggests the probable existence of differences in strength and direction of selection pressures even on very small spatial scales. This is consistent with other studies employing replicated paired sample designs [14,56–58], highlighting the complexity of selection drivers and the difficulties to identify them in natural experimental settings.

## Conclusion

Our findings are in line with recent studies that identified loci under divergent selection between stands growing in contrasting environmental conditions on the local scale in long-lived forest trees [17–19,78]. The increasing number of available genetic markers, also in non-model species, will improve the statistical power to detect such patterns on local scales. Understanding how microenvironmental heterogeneity shapes and maintains the functional genetic variation is especially relevant as this local scale variation is at the base of the population response to future climate. The importance of genetic variation within populations and the strength of selection on small spatial scales have probably been underestimated so far. Especially with respect to climate change, the knowledge about genetic variation and processes that shape the genetic structure on different geographic scales are of utmost importance to develop suitable forest tree conservation and management strategies. Forest management, for instance, could be used to foster natural standing genetic variation and hence *in situ* evolution [79] potentially making unnecessary the use of assisted gene flow or migration.

## Data accessibility

GPS coordinates and SNP genotypes of all individuals included in this study are available on Zenodo public database: https://doi.org/10.5281/zenodo.6345964. The R-script ‘paired_GEA.R’ for analyzing population allele frequencies in paired stands with a known environmental contrast can be found at https://gitlabext.wsl.ch/rellstab/genotype-environment-associations.

## Acknowledgements

We thank Ana Hernández-Serrano, Fernando del Caño, Delphine Grivet, Mario Zabal-Aguirre, Natalia Vizcaíno-Palomar, Enrique Sáez-Laguna, Diana Turrión and Miguel Navarro for help in the field and Carmen García-Barriga (INIA-CIFOR) for assistance in the laboratory. The present study was funded by projects from the Spanish Ministry of Science and Innovation (MICINN): VaMPiro (CGL2008-05289-C02-01/02), LinkTree (EUI2008-03713, under the ERAnet-BiodivERsA call), and TrEvol (CGL2012-39938-C02-00/01), and the IdEx Bordeaux (EcoGenPin, Chaires d’installation 2015). KBB acknowledges an exchange grant of the European Science Foundation (ESF) within the framework of ‘Conservation Genomics: Amalgamation of Conservation Genetics and Ecological and Evolutionary Genomics’ (ConGenOmics).

## Author contributions

MH, JGP, MV and SGCM designed the study; KBB performed the study, did most data analyses and wrote the first draft of the manuscript; FG and CR provided support for data analyses; SGCM contributed to manuscript writing. All authors critically read and revised prior versions of the manuscript.

## Supplementary Material

### S1 Phenotypes

**Fig. S1.1** Boxplots of height (a) and diameter at breast height (DBH, b) for 150 *Pinus pinaster* trees in three stand pairs contrasting north- (N1, N2, N3) and south-facing slopes (S1,S2, S3).

**Table S1.1** Two sample *t*-tests assessing differences in height and diameter at breast height (DBH) of *Pinus pinaster* trees in each pair of north- and south-facing slopes.

### S2 Genetic structure

**Table S2.1** Pairwise differentiation between all six *Pinus pinaster* study stands (comprising north-[N1, N2, N3] and south-facing slopes [S1,S2, S3]) based on 5,024 SNPs. All *F*_ST_ values are significant with *P*< 0.001.

**Fig. S2.1** Principal component analysis (PCA) based on 5,024 single nucleotide polymorphism markers of all *Pinus pinaster* samples from eastern Spain representing three south-facing (S) and three north-facing (N) slopes in a paired sampling design. Each stand is depicted with a different colour and the stand centroid is labelled with the site identifier.

**Fig. S2.2** Fine-scale spatial genetic structure (SGS), plotted as average pairwise Loiselle kinship coefficient against the geographic distance between *Pinus pinaster* trees within pairs of north and south-facing slopes. SGS was strongest for pair N3/S3. *Sp*, intensity of the SGS; ***, significance level of regression slope *P* < 0.001.

### S3 *F*_ST_ outlier detection in south-vs. north-facing slopes

**Table S3.1 (provided as additional spreadsheet file)** Results summary and annotation details of significant single nucleotide polymorphisms detected by the hierarchical models in Arlequin and BayeScanH between *Pinus pinaster* stand pairs of north- and south-facing slopes.

**Fig. S3.1** Plots showing differences in allele frequency between south- and north-facing slopes in all three *Pinus pinaster* stand pairs (*left side*: a, c, e, f, g, i) for the five candidate loci jointly identified by Arlequin and BayeScanH. Pairwise *F*_ST_ between stands is also shown (*right side*: b, d, f, h, j) for each of the five loci.

