## Supplementary material for "Divergent selection in Mediterranean pine stands on local spatial scales"

*Running title:*

Divergent selection in a Mediterranean pine

### S1 Phenotypes

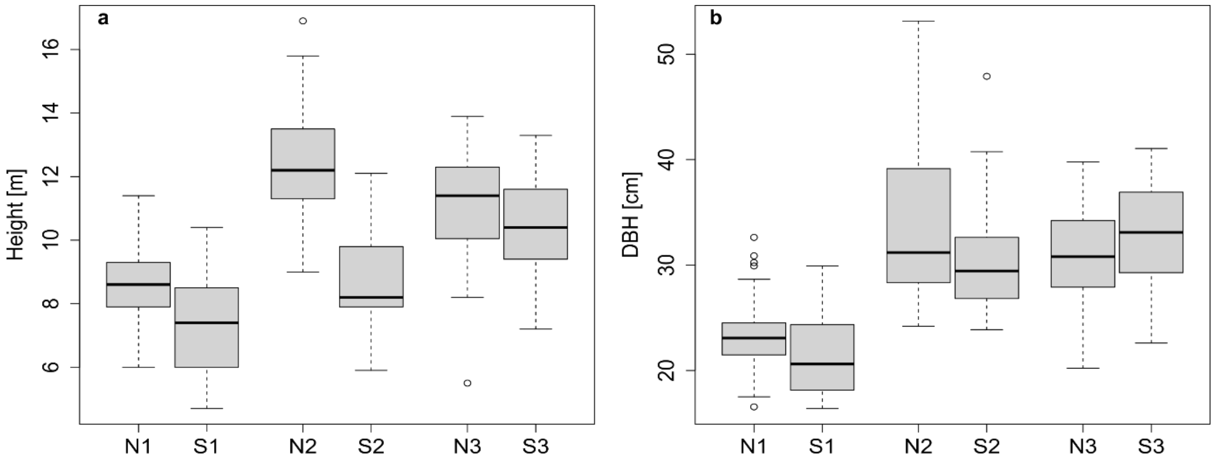


**Fig. S1.1** Boxplots of height (a) and diameter at breast height (DBH, b) for 150 *Pinus pinaster* trees in three stand pairs contrasting north- (N1, N2, N3) and south-facing slopes (S1,S2, S3).

**Table S1.1** Two sample *t*-tests assessing differences in height and diameter at breast height (DBH) of *Pinus pinaster* trees in each pair of north- and south-facing slopes.

| | DBH | | height | |
|---|---|---|---|---|
| | *t* | p-value | *t* | p-value |
| S1/N1 | -1.9861 | 0.0531 | -3.5344 | 0.0009 |
| S2/N2 | -1.6458 | 0.1069 | -7.2921 | <0.0001 |
| S3/N3 | 1.2619 | 0.2135 | -1.0460 | 0.3009 |

### S2 Genetic structure

**Table S2.1** Pairwise differentiation between all six *Pinus pinaster* study stands (comprising north- [N1, N2, N3] and south-facing slopes [S1,S2, S3]) based on 5,024 SNPs. All $F_{ST}$ values are significant with $P < 0.001$.

| | N1 | N2 | N3 | S1 | S2 | S3 |
|---|---|---|---|---|---|---|
| N1 | | 0.0129 | 0.0287 | 0.0041 | 0.0202 | 0.0220 |
| N2 | | | 0.0212 | 0.0165 | 0.0072 | 0.0197 |
| N3 | | | | 0.0331 | 0.0254 | 0.0278 |
| S1 | | | | | 0.0223 | 0.0271 |
| S2 | | | | | | 0.0213 |

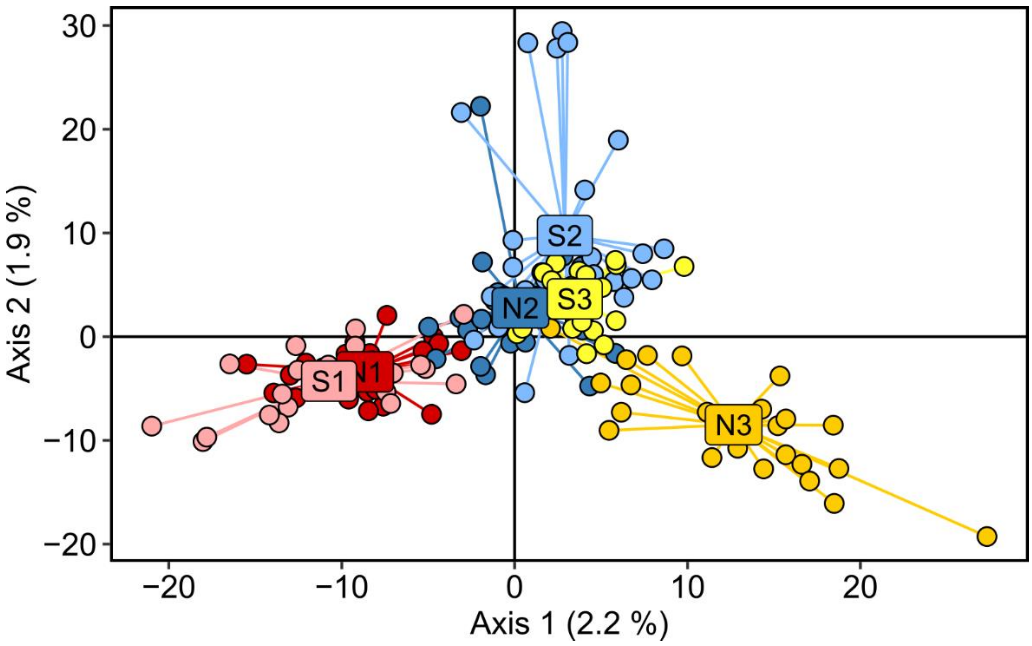


**Fig. S2.1** Principal component analysis (PCA) based on 5,024 single nucleotide polymorphism markers of all *Pinus pinaster* samples from eastern Spain representing three south-facing (S) and three north-facing (N) slopes in a paired sampling design. Each stand is depicted with a different colour and the stand centroid is labelled with the site identifier.

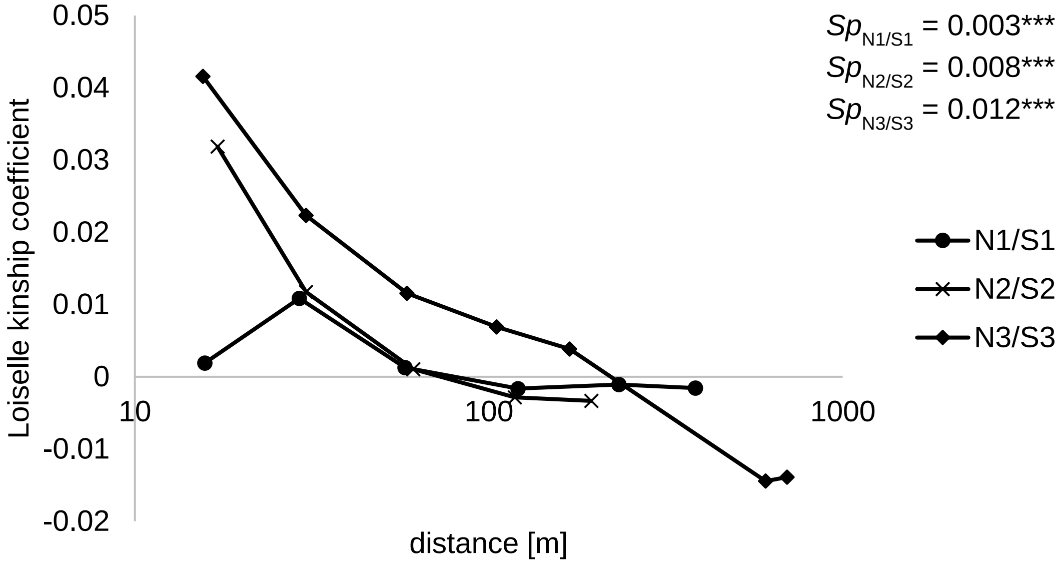


**Fig. S2.2** Fine-scale spatial genetic structure (SGS), plotted as average pairwise Loiselle kinship coefficient against the geographic distance between *Pinus pinaster* trees within pairs of north and south-facing slopes. SGS was strongest for pair N3/S3. *Sp*, intensity of the SGS; ***, significance level of regression slope $P < 0.001$.

### S3 $F_{ST}$ outlier detection in north- vs. south-facing slopes

**Table S3.1 (additional spreadsheet file)** Results summary and annotation details of significant single nucleotide polymorphisms detected by the hierarchical models in Arlequin and BayeScanH between *Pinus pinaster* stand pairs of north- and south-facing slopes.

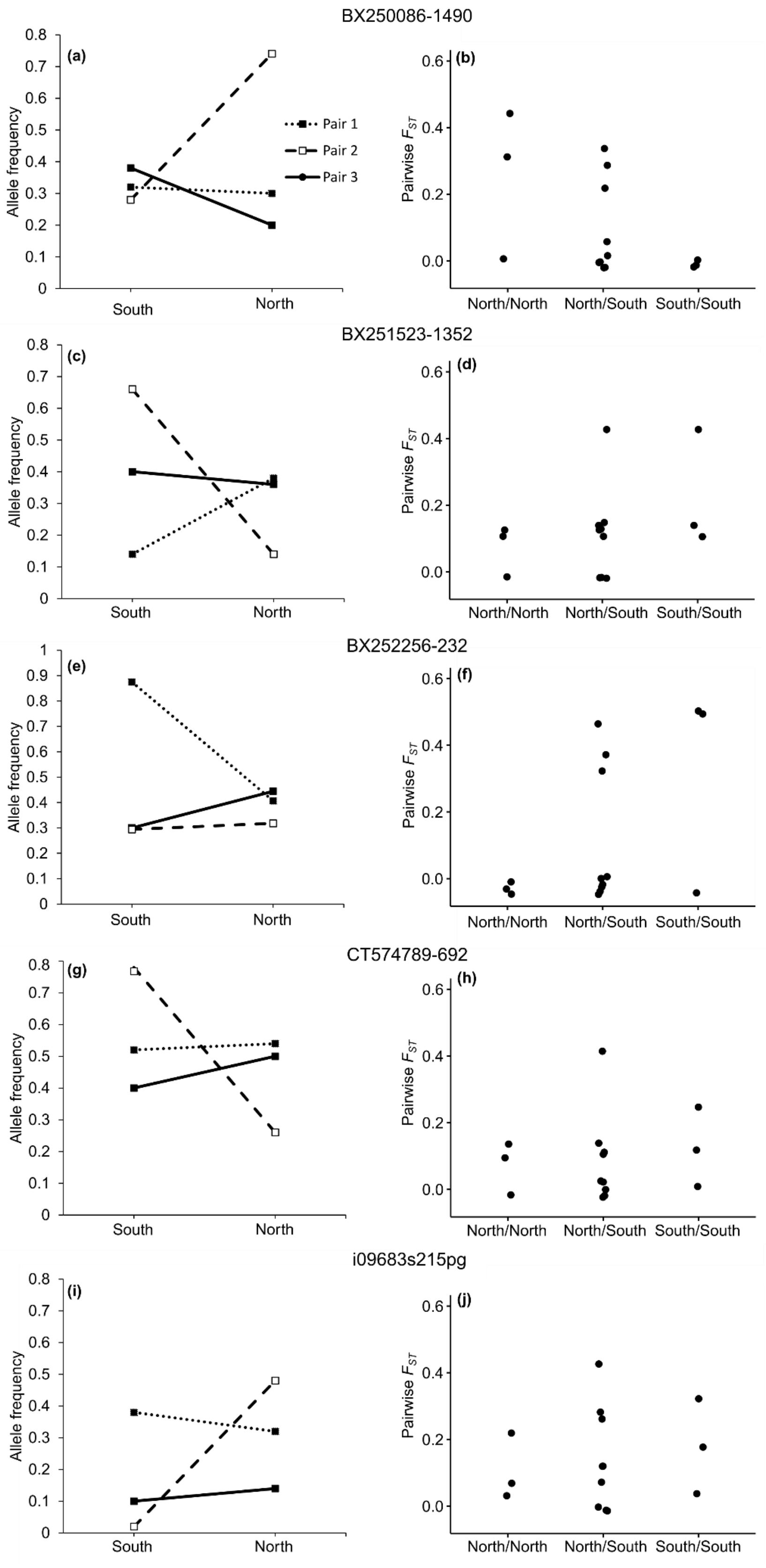


**Fig. S3.1** Plots showing differences in allele frequency between south- and north-facing slopes in all three *Pinus pinaster* stand pairs (*left side*: a, c, e, f, g, i) for the five candidate loci jointly identified by Arlequin and BayeScanH. Pairwise $F_{ST}$ between stands is also shown (*right side*: b, d, f, h, j) for each of the five loci.
